# Integrative Analysis of Spatial and Single-Cell Transcriptomics Reveals Principles of Tissue Organization and Intercellular Communication in Mouse Olfactory Bulb

**DOI:** 10.1101/2020.09.09.290064

**Authors:** Francisco Jose Grisanti Canozo, Zhen Zuo, James F. Martin, Md. Abul Hassan Samee

## Abstract

Intercellular communication and spatial organization of cells are two critical aspects of a tissue’s function. Understanding these aspects requires integrating data from single-cell RNA-Seq (scRNA-seq) and spatial transcriptomics (ST), the two cutting edge technologies that offer complementary insights into tissue composition, architecture, and function. Integrating these data types is non-trivial since they differ widely in the number of profiled genes and often do not share marker genes for given cell-types. We developed STANN, a neural network model that overcomes these methodological challenges. Given ST and scRNA-seq data of a tissue, STANN models cell-types in the scRNA-seq dataset from the genes that are profiled by both ST and scRNA-seq. The trained STANN model then assigns cell-types to the ST dataset. We apply STANN to assign cell-types in a recent ST dataset (SeqFISH+) of mouse olfactory bulb (MOB). Our analysis of STANN’s assigned cell-types revealed principles of tissue architecture and intercellular communication at unprecedented detail. We find that cell-type compositions are disproportionate in the tissue, yet their relative proportions are spatially consistent within individual morphological layers. Surprisingly, within a morphological layer, there is a high spatial variation in cell-type colocalization patterns and intercellular communication mechanisms. Our analysis suggests that spatially localized gene regulatory networks may account for such variability in intercellular communication mechanisms.

## Introduction

All organs are composed of a complex mixture of cells that are further organized into tissues that make up an organ. Organ physiology and function depend on the coordinated action of cells embedded within the tissues of the organ. In pathologic conditions, the normally coordinated organ physiology is disrupted resulting in loss of tissue architecture and eventual organ failure. A mechanistic understanding of a tissue depends on its cell-types and their transcriptional properties, spatial organization, and communication^1^. Methodological advances to address these mechanistic details are essential for the study of development, physiology, and disease^2–8^. Spatial transcriptomics (ST) and single-cell RNA-seq (scRNA-seq) are complementary technologies, each with its own strengths and weaknesses, for studying complex tissues at a single-cell resolution^2^. While scRNA-seq can profile the transcriptional expression of thousands of genes per cell (typically 10-20k)^9^ and characterizes a broad panel of cell-types, it does not preserve the spatial context of cells within the tissue^10^. In order to uncover the spatially varying principles of tissue architecture and intercellular communication, alternative strategies that maintain spatial information are required^11^. The ST technologies complement scRNA-seq by preserving individual cells’ spatial location in their native tissue^10^. However, current ST protocols profile the transcriptional expression of only about half as many genes as scRNA-seq (typically ~1k-10k)^10^, an issue that can make it problematic to identify cell-types in ST datasets. In particular, when the marker genes of different cell-types are absent in an ST dataset, it is unclear if one can assign correct types to the cells in that dataset. Errors in cell-type assignment, in turn, may lead to inaccurate biological conclusions from an ST data analysis.

To harness the potentials of both scRNA-seq and ST, we developed STANN (<u>S</u>patial <u>T</u>ranscriptomics Cell Types <u>A</u>ssignment Using <u>N</u>eural <u>N</u>etworks) -- a neural network that learns from an scRNA-seq dataset to assign cell-types to the cells of an ST dataset (Fig. 1A,B). The inputs to STANN are the transcriptional expression of genes that are shared between the two datasets and an assignment of cell-types to the scRNA-seq cells. Given this information, STANN first learns the mapping of cell-types to scRNA-seq cells (Fig. 1A). STANN next predicts the cell-type of each cell in the ST dataset given the transcriptional expression of the shared genes in that cell (Fig. 1B). We applied STANN to learn cell-type mapping from an scRNA-seq dataset of mouse olfactory bulb (MOB)^12^ and assign cell-types to a recent ST dataset (sequential Fluorescence In Situ Hybridization plus; seqFISH+^13^) of MOB. Utilizing these annotated cell-types, we pursued a detailed investigation of the architecture and intercellular communication mechanisms in MOB (Fig. 1C).

**Figure 1.**
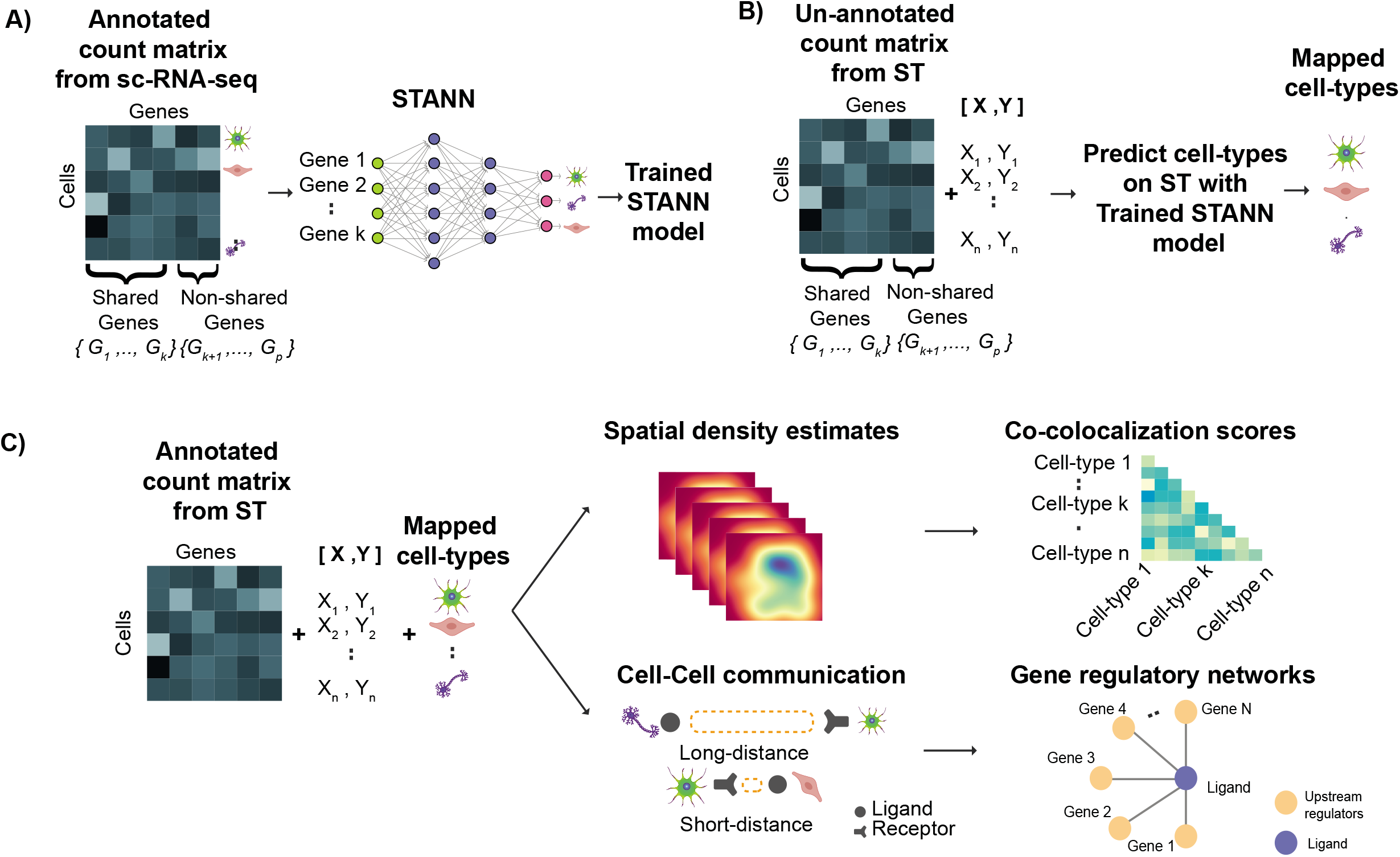
A) Learning stage of STANN, model training on annotated scRNA-seq data. B) ST Cell-type mapping with trained STANN model. C) Downstream analyses on cell-type annotated ST, spatial density maps of locations of cell-types, co-localization scores, intercellular communication and gene regulatory networks.

Our analysis revealed that the proportions of cell-types vary across different morphological layers of MOB, but remain consistent within a given morphological layer. To investigate the spatial organization of different cell-types, we developed methods akin to colocalization analysis in microscopy images^14^. Interestingly, these methods revealed that colocalization patterns between a pair of cell-types vary significantly across different spatial locations of the MOB and even within a morphological layer. This variation ranges from spatially separated occurrences to high degrees of colocalization between the two cell-types. We also find a remarkable variation in intercellular communication mechanisms between the same pair of cell-types, defined by significant co-expression of receptor-ligand pairs^15^, across different locations within a morphological layer. Importantly, by using different sets of receptor-ligand pairs at different spatial locations, a given pair of cell-types also activates different sets of genes at those locations. This suggests that even when cell-type identities do not change, the overall transcriptional signatures of the corresponding cells can change across the tissue giving rise to spatially localized gene regulatory networks.

In summary, the STANN model and the downstream analyses pinpointed the consistent and the variable aspects of the architecture and intercellular communication mechanisms in MOB in unprecedented detail. To our knowledge, this is the first such systematic and rigorous attempt to delineate the principles of tissue architecture and intercellular communication by harnessing the unique features of the ST and scRNA-seq technologies.

## Results

### STANN: a Deep Neural Network Model for the Supervised Mapping of Cell-Types from scRNA-seq to ST Data

Deep neural networks have recently shown remarkable performance in classification tasks^16^. The models can efficiently deal with high dimensional data and learn sufficiently complex, non-linear functions in a data-driven manner to map data points to their respective classes. With proper statistical measures, one can identify the architecture and the parameters that yield an accurate and generalizable model^17^. The STANN (<u>S</u>patial <u>T</u>ranscriptomics Cell Types <u>A</u>ssignment Using <u>N</u>eural <u>N</u>etworks) model in this work implements a fully connected deep neural network (Fig. 1A). The inputs to the model are gene expression values of a cell (input layer) and the outputs are the probabilities of the cell belonging to different cell-types (output layer). Each intermediate layer comprises neurons that linearly combine inputs from the neurons of the previous layer and propagate the result through a nonlinear activation function to the next layer (Methods). The linear weights in each neuron are optimized using the backpropagation algorithm^18^.

We trained STANN on 15 cell-types with 11744 cells and 9031 genes. In order to identify an accurate and generalizable model, STANN performed a hyper-parameter search for the optimal number of intermediate layers, numbers of neurons in each layer, and the activation function for neuron’s in each layer. This identified an optimal model with two hidden layers (with 160 and 145 neurons) and the hyperbolic tangent function as the nonlinear function. In order to test the generalizability of STANN on unseen data, we performed a 10-fold cross validation. On each fold, the data is split into 80% training and 20% testing. Over the 10 folds, the model showed an average accuracy of 99.5 ± 0.1% on the training data and 95.4% ± 0.5% on the separately held-out test data.

STANN also achieved remarkable accuracy for individual cell-types, even for cell-types with small numbers of cells (Fig. 2A). We attribute this capability of STANN to its use of a class-balance aware loss function, that penalizes under or over-represented cell-types. Other current methods do not use this approach^19^; hence it is likely that those methods will be biased to the majority classes, underfit rare cell-types, and their reported performances are likely inflated (Methods).

**Figure 2.**
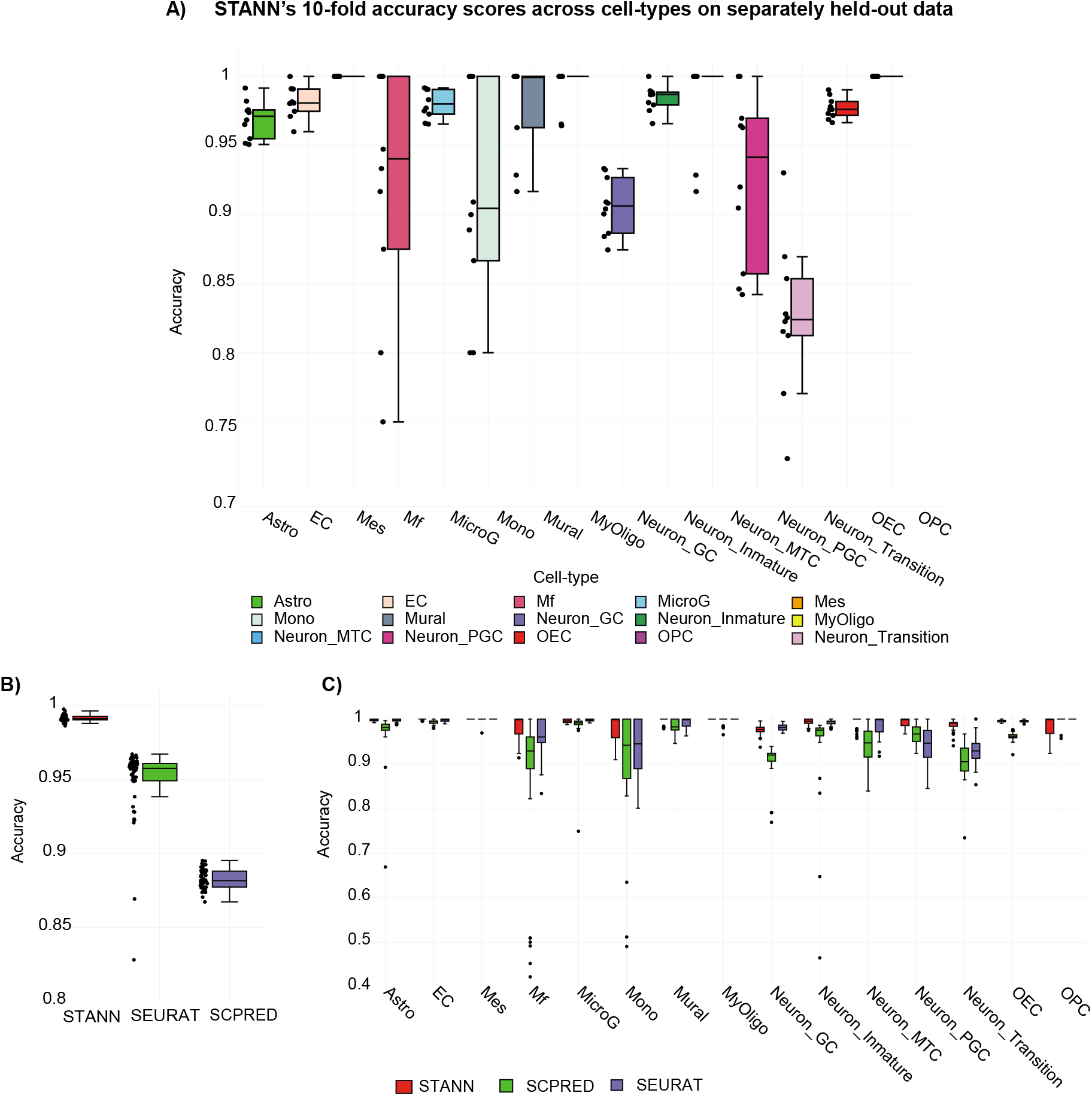
A) STANN’s 10-fold accuracy scores across cell-types on separately held-out data, mean accuracy across all classes is 95.3%. Astro: Astrocytes, EC: Endothelial Cells, OECs: Olfactory Ensheathing Cells, MicroG : Microglia Cells, Neuron_Inmature: Neuron Immature Cells, Neuron_Mature: Neuron Mature Cells, Neuron_GC: Neuron Granule Cells, MyOligo: Myelinating-Oligodendrocyte Cells, Mes: Mesenchymal Cells, Mural: Mural Cells, Mono: Monocytes Cells, Neuron_MTC: Neuron M/T cells, Mf: Macrophages, OPC: Oligodendrocyte Progenitor Cells, Neuron_PGC: Periglomerular Cells. Boxplots showing overall accuracy (B) and cell type specific accuracies for STANN, Seurat’s CCA and scPred (C). Horizontal lines in the boxplots mean the following: center line, median; box limits, upper and lower quartiles; whiskers, 1.5x interquartile range; points, outliers.

To compare STANN with other available tools, we noted that the majority of these methods rely on marker genes to assign cell-types in the ST dataset ^20–22^. However, the MOB scRNA-seq and SeqFISH+ datasets share only 33% of the marker genes (Supplementary Table S1), which makes it unlikely that marker gene based methods will be effective in this case. We therefore used the widely used Seurat method^23^, which first aligns the two datasets in low dimensions before utilizing the marker genes. STANN performed better than Seurat by a margin of 4.9% in our benchmarking runs (Fig. 2B, Methods). We also compared STANN against a scPred^24^, another robust method that combines unbiased feature selection from a reduced-dimension space with support vector machines. STANN performed better than scPred by 10.9% in our benchmarking runs (Fig. 2B, Methods). Briefly, we compared Seurat and scPred against STANN over 50 independent runs. In each run, we simulated an ST dataset by taking a random subsample of ~9k genes (equal to the number of shared genes between scRNA-seq and ST datasets) from the original scRNA-seq data and tested each method on this subsampled data.

We found that STANN outperformed Seurat and scPred in all 50 runs, with accuracies of 99.2%± 0.21% vs. Seurat’s 94.4% ± 4.51% and scPred’s 88% ± 0.64% (Fig. 2B). STANN’s performance was consistently high in rare cell types (<30 cells). For example, for monocytes (Mono), macrophages (MF), oligodendrocyte progenitor cells (OPC), and mural cells, STANN showed improvements of 0.4 – 17% over Seurat and scPred (Fig 2C).

### STANN Provides a Spatially Resolved Annotation of Cell-Types in the Mouse Olfactory Bulb

The mouse olfactory bulb (MOB) SeqFISH+ dataset^13^ comprises six fields of view (FOV) from a slice of adult MOB (Fig. 3A). Each FOV is approximately 0.2 mm X 0.2 mm in size and captures the expression of 10,000 genes from 2050 cells^13^ (Fig. 3A). The MOB is structured into several layers, known as the MOB morphological layers. In a coronal image of MOB, these layers appear in the following order from the center to the outside: granule cell layer (GCL), internal plexiform layer (IPL), mitral cell layer (MCL), external plexiform layer (EPL), and glomerular layer (GL). The FOVs in the SeqFISH+ data span all of these morphological layers^13^ (Fig. 3A; Supplementary Table S2). This dataset has 9031 genes in common with our previously published scRNA-seq data^12^ of MOB. In the rest of this manuscript, we refer to these genes as the “shared” genes. Importantly, the shared genes include only 33% of the marker genes for MOB cell-types (Supplementary Table S1) – suggesting that marker gene based methods^20–22^ would not be ideal for predicting cell-types in this SeqFISH+ data.

**Figure 3.**
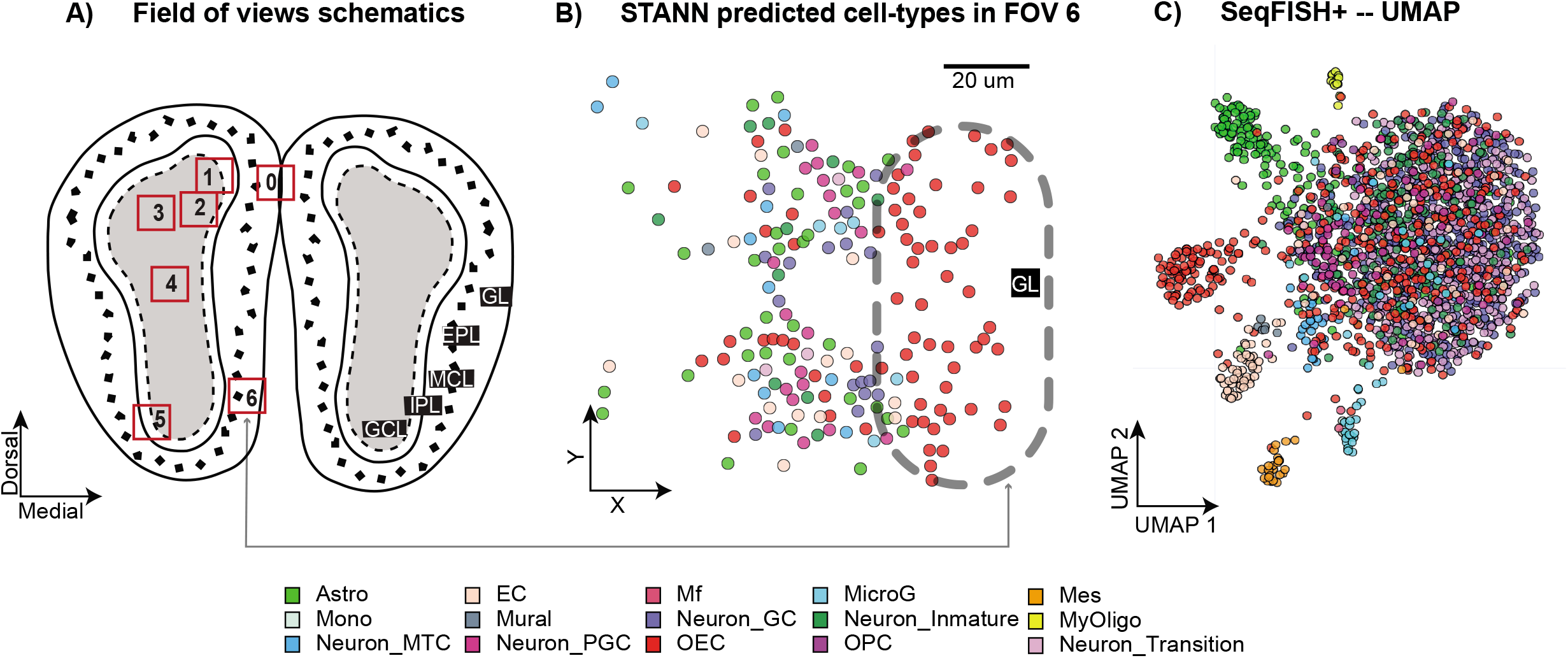
A) A schematic showing the locations of the six fields of views (FOVs) in the MOB SeqFISH+ data. B) Spatial distribution of predictions in FOV six, located in the glomerular layer and external plexiform layer (cf. Fig 4F in Eng. et al^13^). C) model predictions mapped to the uniform manifold approximation and projection (UMAP) embedding of SeqFISH+ data.

As discussed above, we trained STANN on our scRNA-seq dataset^12^ and utilized the trained model to assign cell-types to the SeqFISH+ data. From the original scRNA-seq dataset^12^, 14 major cell-types were found in the WT condition (Supplementary Table S3; Fig. S1). Consistent with previous studies^4,25^, STANN showed an abundance of olfactory ensheathing cells in FOVs originating from the GL (Fig. 3B). STANN also predicted neuronal granule cells as the most abundant cell-type in these six FOVs. This is expected since the majority of the FOVs are located within the GCL or map to an extension of it (Fig. 4A). Overall, STANN revealed a high imbalance in the proportions of cell-types in SeqFISH+ data (all FOVs considered together; Fig. 4B). This likely reflects MOB biology and is not an artifact of STANN, since the MOB scRNA-seq data has a similar imbalance of cell-type proportions (Fig. 4A). However, STANN found subtle differences between the two datasets in terms of the relative abundance of cell-types. For example, the most abundant cell-type across the six FOVs of SeqFISH+ data is neuronal granule cells, which is the second most abundant cell-type in the scRNA-seq data. Similarly, the most abundant cell-type in scRNA-seq, olfactory ensheathing cells, is the third-most abundant cell-type in SeqFISH+. These variations are expected since the SeqFISH+ FOVs constitute a small sample of the scRNA-seq data. Importantly, these variations imply that STANN did not overfit the scRNA-seq data.

**Figure 4.**
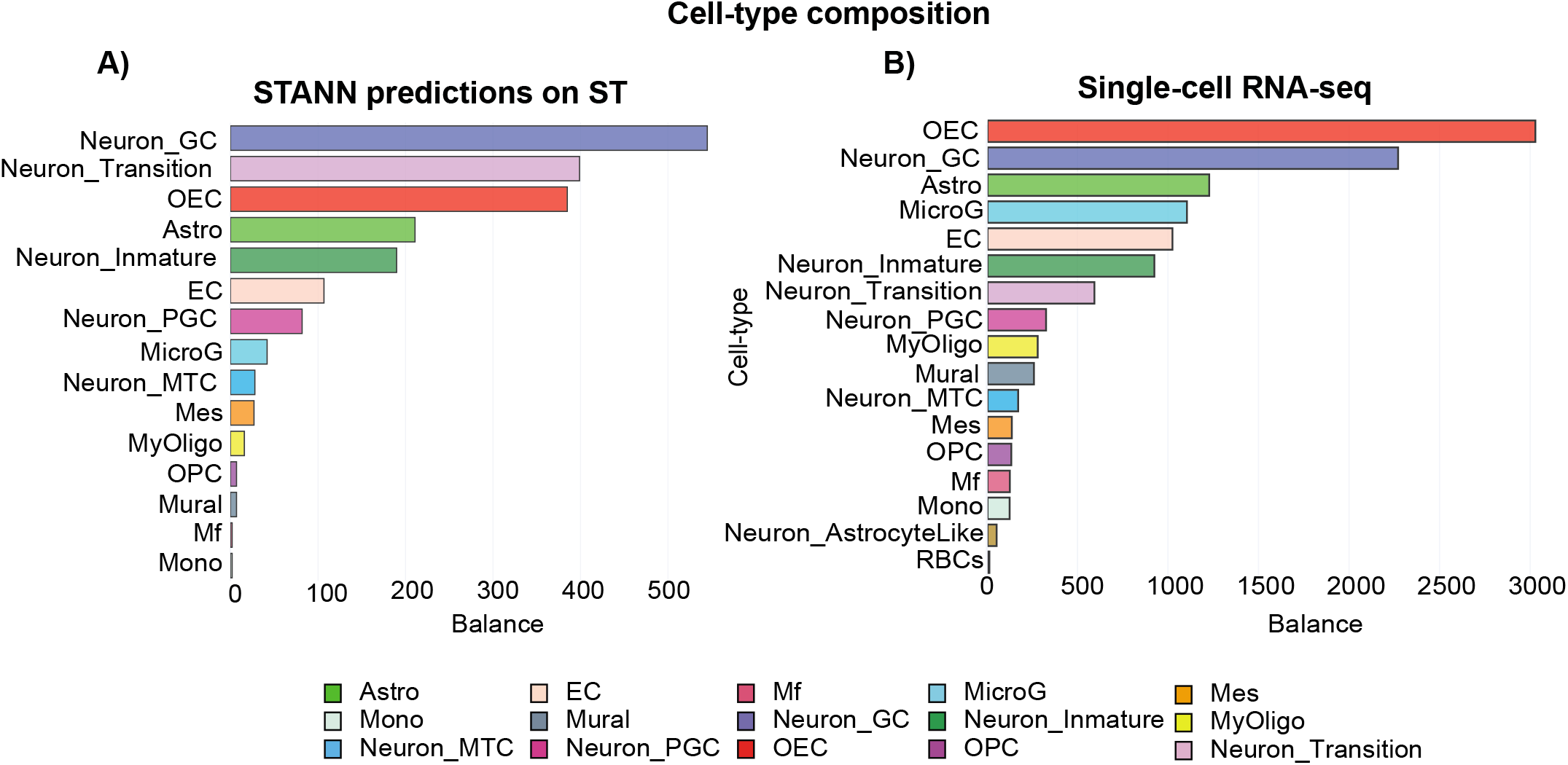
MOB cell-type compositions according to STANN’s predictions (A) and the scRNA-seq data (B).

STANN also highlighted the importance of utilizing models to learn cell-types from shared genes. In particular, the SeqFISH+ data UMAP (Uniform Manifold Approximation and Projection; Fig. 3C) built from the shared genes did not converge to a good separation of neuron clusters, implying that the shared genes do not capture sufficient variation to have a straightforward classification of those cell-types. However, as STANN shows, it may be possible for nonlinear functions to integrate information from the shared genes and accurately classify all cell-types.

### STANN Predictions Reveal Consistency in Cell-Type Composition but Variability in Colocalization Patterns across the Tissue

STANN predictions revealed that the proportion of cell-types are consistent within a given morphological layer of MOB, but may vary between two different morphological layers. Interestingly, the patterns of colocalization between a pair of cell-types can vary substantially even within a given morphological layer.

We define the cell-type composition of an FOV as the proportion of different cell-types in it.

Within a morphological layer, STANN revealed that the cell-type compositions of each FOV are very similar (based on both Chi-square test and relative entropy; Methods; Supplementary Table S4). For example, the three FOVs located within the GCL have nearly identical cell-type compositions (FOVs 2, 3, and 4 in Fig. 5A; Supplementary Table S4). However, FOVs differ in cell-type compositions when they span different morphological layers. For example, FOV 1 (spanning GCL and IPL) and FOV 5 (spanning GCL, IPL, EPL and MPL) have significantly different cell-type compositions (Fig. S2, Supplementary Table S4). The same applies for the FOVs 0 (spanning GL and EPL) and 4 (spanning GCL).

**Figure 5.**
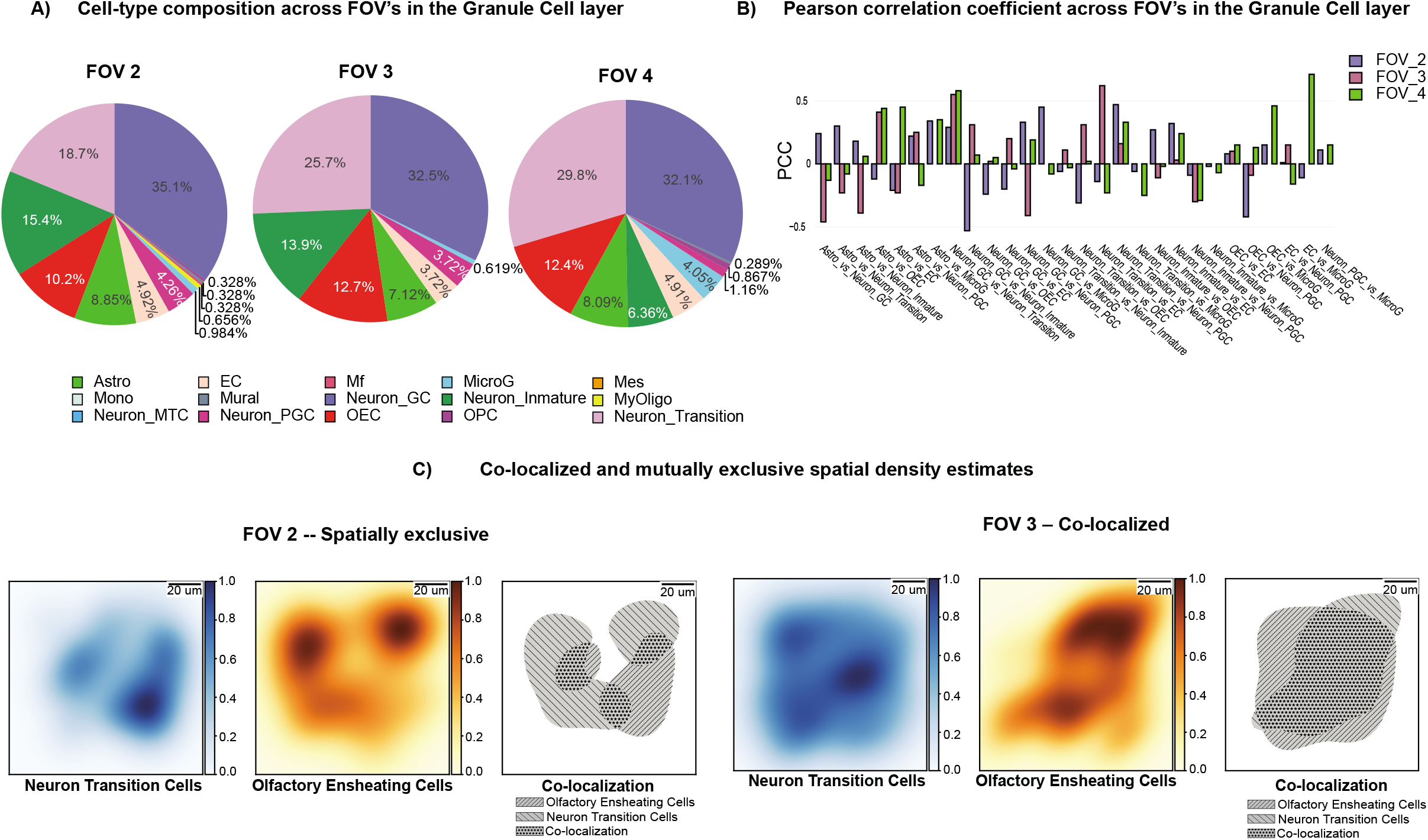
a) Composition of cell-types across different FOVs in the Granule Cell Layer (GCL). b) PCC scores of FOVs that mapped the GCL c) Kernel density maps of Neuronal Transition cells vs Olfactory Ensheathing cells in two different FOVs located within the GCL layer, showing completely opposite co-colocalization patterns.STANN Predictions Reveal Spatially Localized Intercellular Communication Networks

We next quantified colocalization between any pair of cell-types in a given FOV. For each cell-type predicted by STANN in an FOV, we calculated a multivariate kernel density^26^ map from its locations within the FOV (Fig. 5C; Methods). We computed the Pearson correlation coefficient between the probabilities of observing the two cell-types across all spatial coordinates in that FOV. This colocalization analysis, built upon previous studies of colocalized fluorescence probes^27^, identified cell-type pairs with significant colocalization (strong positive correlation) or cell-type pairs that were spatially separate (strong negative correlation) in each FOV (Fig. 5B,C). Interestingly, we found a high variation in colocalization patterns between every pair of cell-types across the FOVs (Fig. S3). The three FOVs located within the GCL (FOV 2, 3, and 4) demonstrate this point. The most conservative prediction will be that FOVs that share the same morphological layer or are adjacent, will be more similar than FOVs located in different morphological layers or at a distance. While this conservative prediction holds for 36% of cell-type pairs in FOVs 2, 3, and 4, we found that 64% of the cell-type pairs in these FOVs show highly variable colocalization patterns, ranging from spatial separation to significant colocalization across the FOVs (Figure 5B).

Intercellular communication is critical for tissue function^28–33^ and is also associated with disease states^21,34,35^. Identifying the communicating pairs of cell-types and their communication mechanisms (in terms of receptors and ligands) is currently a major focus of scRNA-seq studies. Recent studies suggest that the communications between certain cell-types may be enriched within specific parts of a tissue (commonly termed microenvironments or niches^34,36^). However, to what extent and how intercellular communication varies across a tissue is still an open question^15^. It is also unclear if the communication between two cell-types changes with their abundance or colocalization in different parts of a tissue.

We applied CellPhoneDB^15^, an intercellular communication analysis tool, on STANN’s predicted cell-types in each FOV of the SeqFISH+ data (Methods). For every pair of cell-types in an FOV, CellPhoneDB identifies the receptors and ligands that are significantly and specifically enriched between those cell-types. However, CellPhoneDB does not take the spatial location of cell-types into account and could incorrectly infer communication between two spatially separated cell-types, *e.g.,* when they each communicate with a third cell-type but not with each other (Fig. 6A). To make our analysis robust against this type of incorrect inferences, we have applied an additional filtering on the intercellular communications between spatially separated cell-types (Methods). This eliminated 63.07% of the communications inferred by CellPhoneDB. Although we were over-conservative in this step, the possibility of such a large fraction of inferences being incorrect underscores the importance of studying intercellular communication with spatial information.

**Figure 6.**
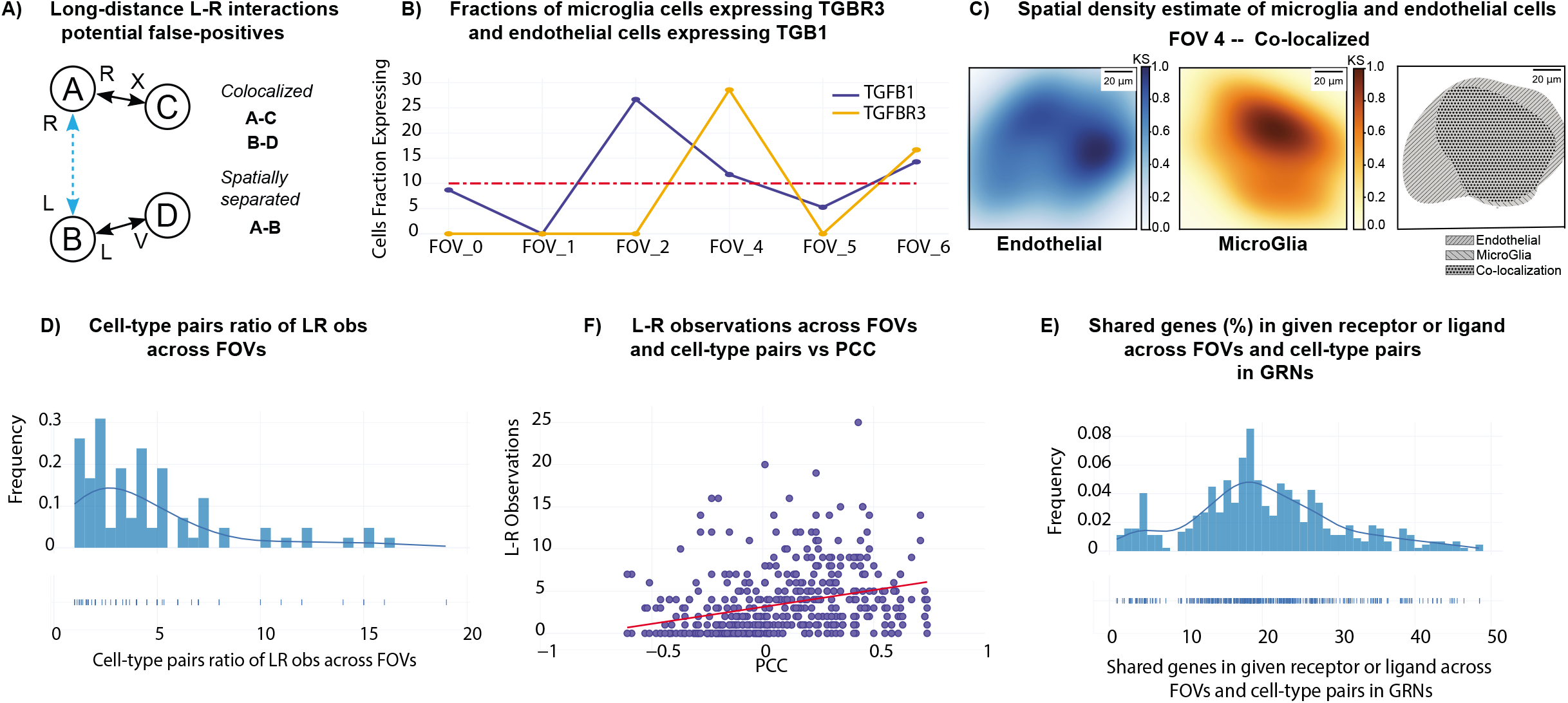
A) Outline of the scheme to identify potential false-positive long-range ligand-receptor communications. B) Variability in expression of TGF β ligand in microglia cells and type III TGF β receptor in endothelial cells. Vertical axis represents a fraction of the cell-type population that expresses the ligand/receptor and the horizontal axis represents FOV (Field of View). C) Kernel density maps of endothelial cells and microglia, and their co-colocalization pattern in FOV 4 (located within the GCL layer). D) Ratio between the maximum and the minimum numbers of ligand-receptor pairs utilized by each pair of cell-types across FOVs. E) Number of ligand-receptor observations across FOVs and cell-type pairs vs Pearson Correlation Coefficient (PCC) F) Shared genes in receptor or ligand GRNs across FOVs and cell-types pairs.

This analysis discovered widespread intercellular communication in MOB while capturing bona fide receptors and ligands. For example, we found endothelial and microglial cells communicating through the Type 3 TGF-β receptor and the Type 1 TGF-β ligand in FOV 4 (Fig. 6B). Previous studies have experimentally validated this TGF family receptor-ligand pair in MOB^13,37^. These two cell-types are also colocalized in this FOV (Pearson correlation coefficient for colocalization = 0.71) (Fig. 6C), further supporting the possibility of their communication. Similarly, we found neuronal transition cells and olfactory ensheathing cells to communicate through the BMP pathway ligand Bmp7 and the receptor Slamf1 in FOV 6, although the two cell-types are spatially separated in this FOV (Pearson correlation coefficient for colocalization = −0.42). The BMP pathway is classically known to mediate long-range intercellular communication^38,39^. As expected overall, we found the most abundant receptors and ligands are from the well-known signaling pathways such as Fibroblast growth factor (FGF), Notch, Bone morphogenetic protein (BMP), Transforming growth factor beta (TGF-β), and Wnt ^40–43^.

The analysis also revealed the details of spatial variation in intercellular communication across the FOVs. On average, a given cell-type pair communicates in 4 FOVs (range: 1-7). We observe this variation even when we focus on FOVs from the same morphological layer. For FOVs 2, 3, and 4, which are from the GCL layer, we see that 25% of cell-type pairs communicate in one FOV, 39% in two and 36% in all three FOVs. Some intercellular communications are specific to an FOV or to a particular morphological layer. For example, olfactory ensheathing cells to astrocytes cells communicate only in FOVs 2, 3, and 4 located specifically in the GCL layer. Similarly, myelinating-oligodendrocyte cells and neuronal transitional cells communicate only in FOV 0 that covers the GL, EPL layers. On the other hand, certain pairs of cell-types communicate in nearly every FOV and their communications appear to be a common aspect of MOB. These examples include the intercellular communication between astrocytes and endothelial cells, and neuronal granule cells and endothelial cells.

Interestingly, we found that a cell-type pair often uses different sets of receptors and ligands in different FOvs. On average, the number of receptors and ligands used by two cell-types in different FOVs can change by 3.5-fold (Fig. 6D; ratio of the maximum and the minimum number of receptors-ligands used across different FOVs; range: 0-19), with minimal overlap between the sets of receptors and ligands across FOVs (Median similarity between cell-type pairs in receptor-ligand usage: 3%, range: 0-42%). We observed this variation even across FOVs where the two cell-types colocalize. For example, endothelial and neuron transitional cells colocalize in FOVs 3 and 5 (Pearson Correlation Coefficient > 0.5), but only 7% of their 17 ligand-receptor pairs are common in these FOVs. We found that a higher co-localization between cell-types shows a positive trend with a higher number of ligands and receptors (Fig. 6E), although there are instances when cell-types co-localize but do not communicate.

The above analyses implied spatially variable usage of receptors and ligands for intercellular communication. We next asked if the gene regulatory network (GRN) of a receptor or ligand, *i.e.,* the genes in regulatory relationships with it, in a given cell-type may change across FOVs. This concept of spatially localized GRNs has been noted in the literature^44^. However, to our knowledge, the existence and role of spatially localized GRNs in mediating intercellular communication have never been discussed.

We applied SCENIC^45^, a GRN construction algorithm, on STANN’s predicted cell-types in each FOV (Methods). We found that the GRN of nearly every receptor and ligand in every cell-type changes across FOVs, suggesting that spatially localized GRNs are very common in MOB. For example, endothelial cells use the TGFβ-1 ligand to communicate with microglia cells in FOVs 6 and 4 (Fig. S4), yet only 8.3% of genes in the GRN of TGFβ-1 in endothelial cells are shared in these two FOVs. On average, only 20.38% (Fig. 6F; range 0.93%-48.46%) genes in the GRN of a receptor or ligand in a cell-type were common across FOVs.

Overall, our analyses in this section revealed widespread spatial variation in intercellular communications. A pair of cell-types often interacts in a region-specific manner and may communicate through utilizing very different sets of receptors and ligands. These mechanistic variations are associated with spatially varying gene regulatory networks, pointing to a new level of heterogeneity in single-cell data where cells of the same type show subtle changes in GRNs of receptors and ligands depending on their spatial location.

## Methods

### Cell type annotation in scRNA-seq data

We collected the annotated cell-types from our previous scRNA-seq dataset of mouse olfactory bulb (MOB)^12^. Following the common steps for cell-type annotation in scRNA-seq data^23,46^, we first ranked genes based on the variance-to-mean ratio of their expression values. Then, using principal component analysis, we computed a low dimensional manifold that best explains the variance in the data, and further reduced the dimensionality into two dimensions using uniform manifold approximation and projection (UMAP). We then clustered cells in this two-dimensional space using the Louvain community detection algorithm. Finally, based on marker genes detected using the Wilcoxon rank-sum test, we annotated the clusters (Fig. S1).

### Data normalization

For the scRNA-seq data, Scanpy toolkit 1.3^47^ was used to perform standard pre-processing, including normalization of library-size with 10,000 reads per cell, log transformation, regression of effects of total counts per cell and the percentage of mitochondrial genes expressed. We finally scaled the data to unit variance (clipping values exceeding standard deviation greater than 10).

Following the original SeqFISH+ manuscript, we performed a quantile normalization on the normalized count matrix of each gene in the scRNA-seq and seqFISH+ datasets to correct for cross-platform differences^13,48,49^. Finally, we applied Min-Max scaling to scale the data between 0 and 1. This final step was used to ensure efficient updates in the gradient descent optimization of our neural network model.

### STANN architecture and training

We implemented a multi-layer perceptron model and searched for its optimal architecture (using random initializations in terms of the number of hidden layers, the number of nodes in hidden layers, and the activation functions) using the TensorFlow framework. This approach identified a model with two hidden layers (with 165 and 145 nodes, hyperbolic tangent activation function, and a learning rate of 0.01) as the optimal choice for our preliminary model on the MOB scRNA-seq data. The input layer of this model has size equal to the number of genes that have been profiled under both scRNA-seq and sc-ST. The output layer is a softmax layer with size equal to the number of cell-types detected above from the scRNA-seq data.

The softmax output layer is defined as:

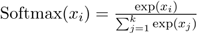

Where all the *x_i_* values are the elements of the output of the fully connected layer immediately preceding it, *k* is the number of classes and 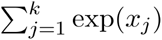 is the normalization term, which ensures that all output will sum to one, and thus constitute a probability distribution.

The hyperbolic tangent function is defined as:

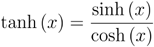

Where tanh (*x*) is the hyperbolic tangent function evaluated at *x.*

We used categorical cross-entropy (CE) as the loss function to train the model. CE is defined as:

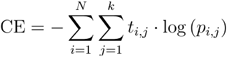

Where *N* is number of observations (data points), *k* is the number of classes, *t_i,j_* is the binary indicator whether class *j* is the correct classification for observation *i*. Finally, *p_i,j_* is the predicted probability that observation *i* belongs to class *j*.

However single-cell datasets are usually imbalanced, so we used an alternative loss function that penalizes under or over-represented cell-types based on their number of cells in the dataset, defined as:

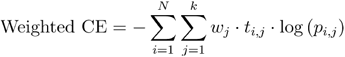

Where *w_j_* is class weight balance defined as:

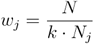

Where *N_j_* is the number of observations in class *j*. In our analysis, this loss function remedied the class-imbalance issue and was our key to achieving superior performance for cell-type assignment^19^. Most of the current methods are not sensitive to class-imbalances. Hence it is likely that those methods will be biased to the majority classes, underfit rare cell-types, and their reported performances will likely be inflated.

### STANN Implementation

STANN is implemented in Python 3 using Keras and its TensorFlow^50^ back end. The model can be trained in a local computer within minutes for a dataset of ~10K cells and ~10K genes. Training on CPU or GPU is supported using Keras and TensorFlow. The hyperparameter optimization is implemented using keras-tuner.

### STANN comparison against alternative methods

We compared Seurat^23^ and scPred^24^ against our model over 50 runs. In each run, we subsampled 2500 cells randomly out of the main scRNA-seq dataset. From this subsampled dataset, we then eliminated all the genes that are not shared between the original scRNA-seq and SeqFISH+ datasets; but we retained the cell-types as computed from the original scRNA-seq dataset. This way, we could treat the subsampled dataset as a simulated ST data where we already know the true cell-types. We evaluated each method on this subsampled dataset through 10-fold cross-validation. We found that Seurat achieved a mean estimated accuracy = 94.4 ± 4.51%, and scPred a mean estimated accuracy = 88.26 ± 0.64%, compared to our method with a mean estimated accuracy = 99.2 ± 0.21% (Fig. 2B).

### Colocalization analysis

The method is based on principles of co-colocalization analysis in microscopy images^27^. Briefly, we first use kernel density estimation to estimate the density of each cell-type in each FOV. Kernel density estimation is a method to estimate the density of a random variable from a finite sample size^26^. For any pair of cell-types, we then used the Pearson’s Correlation Coefficient (PCC)^27^ between their kernel densities in an FOV as a measure of their colocalization in that FOV. Details of the kernel density estimation step is as follows.

The kernel density estimator is defined as^26^.

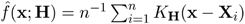

where*K*_H_ is a kernel function that depends on H, the symmetric and positive-definite bandwidth matrix. x is an estimation point and *X_i_* is an observation point. We use the scaled and translated normal kernel function:

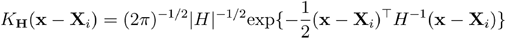

which is a normal density centred at *X_i_* and with variance matrix H. Normal Kernel is the most widely used multivariate kernel function. We used the kernel density estimator from the R package ks^51^. This package estimates H through cross-validation.

We used Pearson’s Correlation Coefficient (PCC)^27^ to measure the similarity between the kernel densities of two cell-types in an FOV. It is the covariance normalized by the product of their standard deviations:

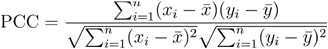

where *x_i_* represents the probability of observing cell-type *x* at the *i^th^* location in that FOV, as estimated in the kernel density for *x.* Similarly, *y_i_* represents the probability of observing cell-type *y* at the *i^th^* location in that FOV, as estimated in the kernel density for y. Here *n* is the number of discrete locations in this FOV. We did not apply PCC on the overall kernel densities, we selected only those locations where probabilities are greater or equal to the 2-quantiles of the kernel densities.

### Ligand-Receptor analysis

With STANN’s predictions of cell-types in each FOV of the SeqFISH+ data, we identified every pair of communicating cell-types and the receptor-ligands that they use. We used the tool CellPhoneDB^15^ for this analysis. CellPhoneDB uses a curated database of receptor-ligand pairs and constrains that a receptor or ligand should be expressed in at least 10% cells of a cell-type to be considered for this analysis. Based on an empirical null distribution, the tool then identifies receptors-ligands that are significantly and specifically enriched within a pair of cell-types.

### Eliminating potential false-positive predictions of long-range intercellular communication

To flag potential false-positive predictions of intercellular communication between spatially separated cell-types, we checked if the predicted ligand-receptor pairs are used for communication with other co-localized cell-types. In particular, we perform the following check for each spatially separated but communicating cell-type pair A-B (PCC < 0). Let the communication between A and B is mediated by the ligand L and the receptor R. We eliminate this ligand-receptor pair from consideration if A is co-localized with cell-type C and communicates with C using L, and if B is co-localized with cell-type D and communicates with D using R. We outline this scheme in Fig. 6A. We reported the remaining ligand-receptors between A-B as potential mediators of long distance communication between A-B (Supplementary Table S5).

### Gene Regulatory Networks

We used pySCENIC^45^ to compute gene regulatory networks for each cell-type within each region of the tissue. Based on these networks, we identified the genes having regulatory relationships with the receptors and ligands mediating intercellular communication.

### Jaccard Similarity coefficient

We used the jaccard similarity coefficient (also known as Jaccard Index) to gauge similarity between two finite sample sets *A* and *B,* e.g., between sets of receptors and ligands across FOVs. The Jaccard index is defined as the size of the intersection (*N*(*A* ⋂ *B*)) divided by the size of the union (*N*(*A* ⋃ *B*)) of the sample sets, it’s expressed as:

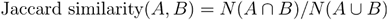

### Statistical comparison of cell-type compositions

To compare cell-type compositions between a pair of FOVs, we used the chi-square test of independence on the counts of different cell-types in those FOVs. We used a p-value cutoff of 0.01 after Bonferroni correction (Supplementary Table S4). We further corroborated these results using relative entropies as follows. Let *x* and *y* denote the compositions of cell-types in the two FOVs. We then computed the KL-divergences of *x* given *y* and also of *y* given *x.* Let *p* and *Q* denote these two relative entropy terms. We checked both the arithmetic and the harmonic means of *p* and *q* and confirmed that the results are in agreement with our chi-square test, i.e., we confirmed that the cases where we could not reject the null hypothesis are indeed the cases with the lowest mean relative entropies.

All analyses were performed using Scipy v1.5.2.

### Mouse Models

Wild-type mice C57BL/6J of post-natal day P40 (male) was used for the olfactory bulb SeqFISH+ experiments^13^. The olfactory bulb scRNA-seq data^12^ was generated from wild-type male and female C57BL/6NJ mice at 14 weeks of age.

## Discussion

Mammalian brains are complex ecosystems of several millions to hundreds of billions of cells^52,53^. Brain cells exhibit a high degree of spatial organization^54,55^ and intercellular communication^2^, which are disrupted in nearly every brain disease. A critical step toward understanding physiology and disease of the brain is to chart the spatial organization of cells and their communications in this tissue.

By integrating scRNA-seq (single-cell transcriptomics) with ST (spatial transcriptomics) data, and performing a series of systematic analyses, we have charted the spatial organization of cell-types and their communications in mouse olfactory bulb (MOB). Our analyses revealed both consistent and variable aspects of cell organization and intercellular communications in the MOB. Interestingly, we found more aspects to be spatially variable than consistent. On the one hand, as we had expected, we found that the proportions of cell-types vary across different morphological layers of MOB, but remain consistent within a given morphological layer. On the other hand, we found remarkable variations in their colocalization patterns and intercellular communication mechanisms throughout MOB. The analysis points to a widespread use of spatially localized gene regulatory networks in the MOB. While the motivation of our study was indeed to discover spatially variable principles of tissue architecture and function, without studying more samples, it is impossible to state which of these variable aspects are characteristic of MOB. For example, we have observed spatially localized gene regulatory networks; however, we cannot comment whether we would observe the same network at the same MOB location across every sample.

The key to this work’s success is a neural network model, STANN, that learns to assign the cell-type to a cell by modeling scRNA-seq data. The trained STANN model can assign cell-types to every cell in an ST data. The two data sources do not profile the same set of genes, so STANN uses only the common genes to both data sources. STANN was highly accurate in this modeling task, but it would be impossible if we had relied only on the well-known marker genes of the MOB cell-types. The ST dataset in this study^13^ shares only 33% of the marker genes with the scRNA-seq dataset^12^, readily implying that the marker gene based methods^20–22^ for integrating different single-cell datasets would not suffice in this case. STANN filled in this gap, and even performed better than the other state of the art methods, such as Seurat^23^ and scPred^24^, that do not rely on marker genes. We attribute STANN’s success primarily to its ability to identify complex nonlinear functions that map the expression values of shared genes in a cell to its cell-type. This highlights that to integrate two single-cell transcriptomic datasets, their shared genes do not necessarily have to include all the marker genes. One can solve this task as long as the shared genes capture sufficient information and the model is sufficiently complex (see below). Another important aspect boosting STANN’s performance was its use of a class imbalance aware loss function. This helped STANN to accurately model rare cell-types, which is very common in single-cell studies. In our comparisons, the widely popular method Seurat lost much of its performance due to rare cell-types.

Although STANN performed very well in integrating the scRNA-seq and ST datasets, it is not guaranteed that such a solution will always exist. From a model design perspective, it is important to determine the cases when it is possible to integrate a given scRNA-seq and an ST dataset. Answering this question also enables us to answer a broader question: which genes should we select in an ST assay if the technology limits the number of genes?

We note that, given a choice of the model (*e.g.,* a support vector machine or a neural network), we can establish a necessary condition on the shared genes that should hold for the model to accurately learn cell-types. Intuitively, for any choice of model, the shared genes need to capture the minimum variance necessary to distinguish the cell-types. Starting with the set of genes in the scRNA-seq dataset ranked according to their variability, one can then train cell-type mapping models (such as STANN) using an increasingly larger number of the most variable genes. The performance of the models in this process should never decrease since each model will use more variance in gene expression than its previous models. Let *R* denote the number of genes used in the first such model that learns cell-type mapping with a satisfactory performance and the performance “saturates” for all models that use more genes.

Then, for the model to be able to correctly learn cell-types using the shared genes, it is a necessary condition that the shared genes have rank ≤*R.* This necessary condition also enables us to find a trade-off between performance in cell-type learning and the number of genes to be profiled in an ST assay. For example, when we do not want to use as many genes as necessary for the model to saturate in performance, we can pick up a sub-optimal performance threshold and decide to profile only the corresponding genes in our ST assay. We tested this condition on the current dataset. Interestingly, we find that only about 5000 genes are necessary to accurately model cell-types in this dataset (Fig. S5). These 5000 genes include only 30.59% of the marker genes, further underscoring the point that with a proper model, we do not need to rely entirely on marker genes for cell-type learning.

The quality and availability of scRNA-seq and ST datasets are increasing at a fast pace, urging for methods to integrate these datasets accurately and without depending on expert-derived prior information, e.g., information on marker genes. The STANN model and the follow-up analyses were an initial step toward this goal. We anticipate that our observations will fuel the development of more sophisticated methods to analyze colocalization patterns and intercellular communication over long and short distances. We anticipate seeing more applications of these tools and methods on datasets collected from developmental stages and diseased tissues. Finally, our work has paved the way toward comparative spatial transcriptomics between healthy and diseased samples. We anticipate future studies will characterize the spatially aberrant transcriptomics of diseased samples enabling the design of targeted therapies.

## Data Availability

The datasets analyzed in this study are available with the respective manuscripts and their supplementary information^12,13^. The accession number for the raw single-cell RNA sequencing data is GEO: GSE121891^12^. We collected the spatial transcriptomics SeqFISH+ data from the authors’ site at https://github.com/CaiGroup/seqFISH-PLUS

## Software Availability

https://github.com/sameelab/STANN

## Acknowledgments

This work was supported by grants from the National Institutes of Health (HL127717, HL130804, HL118761 (J.F.M.); Vivian L. Smith Foundation (J.F.M.), State of Texas funding (J.F.M.), Fondation LeDucq Transatlantic Networks of Excellence in Cardiovascular Research (14CVD01) ‘Defining the genomic topology of atrial fibrillation’ (J.F.M.)

## Supplementary Figure Legends

Figure S1. A) Cellular composition of mouse olfactory bulb visualized using uniform manifold approximation and projection (UMAP). B) Heatmap illustrating the genes most highly enriched in each cluster, with each column representing a gene, and each row representing the average expression level of that gene in each cell type cluster. C) Individual single-cell transcriptomes were colored according to cluster/cell identity.

Figure S2 A) Composition of cell-types across different FOVs and morphological layers

Figure S3 A) Colocalization between cell-types pairs in SeqFISH+ data, quantified as Pearson correlation coefficient (PCC) across different fields of views (FOVs). Horizontal lines in the boxplots mean the following: center line, median; box limits, upper and lower quartiles; whiskers, 1.5x interquartile range; points, outliers

Figure S4. TGF beta ligand upstream regulators in FOV 6 and FOV 4.

Figure S5. A) Metric scores of different models ran with varying numbers of genes/features.

## Supplementary Table Titles

Supplementary Table S1. Marker genes from Tepe et al, 2018 and their intersection with SeqFISH+

Supplementary Table S2. Morphological layer annotation of each Field of Views (FOVs) (also see Figure 3)

Supplementary Table S3. Acronyms of each cell-type referred in this study.

Supplementary Table S4. Statistical comparison of cell-type compositions results

Supplementary Table S5. Ligand-receptors potentially mediating long-range intercellular communication.

